# Cortisol Predicts Antidepressant Treatment Outcome, Memory Improvement, and Brain Response to Negative Emotions: The Importance of Aging

**DOI:** 10.1101/450213

**Authors:** Felipe A. Jain, Colm G. Connolly, Victor I. Reus, Dieter J. Meyerhoff, Tony T. Yang, Synthia H. Mellon, Scott Mackin, Christina M. Hough, Alexandra Morford, Elissa S. Epel, Owen M. Wolkowitz

## Abstract

**Background:** Studies testing the relationship between cortisol levels, depression, and antidepressant treatment response have yielded divergent results suggesting the possibility of moderators of a cortisol effect. Several studies indicate that age may moderate the relationship between cortisol and psychopathology. In patients with Major Depressive Disorder (MDD), we studied the interactive effects of age and cortisol on predicting diagnostic status, improvement in mood and memory function with antidepressant treatment, and brain response to negative emotional stimuli.

**Methods:** 66 unmedicated patients with MDD and 75 matched healthy controls had serum assayed at pre-treatment baseline for cortisol. Logistic regression was used to determine an association of age, cortisol and their interaction with MDD diagnosis. Thirty-four of the MDD participants (age range: 19-65 years; median: 36) underwent treatment with a selective serotonin reuptake inhibitor (SSRl) for 8 weeks. Clinician and self-ratings of depression symptoms, as well as tests of verbal and visual delayed recall were obtained at baseline and post treatment. Moderation analyses determined the effect of age on the relationship between baseline cortisol and treatment outcome. A separate sample of 8 MDD participants prospectively underwent fMRI neuroimaging and cortisol collection while viewing negative emotional faces.

**Results:** Age moderated the effects of cortisol on predicting MDD diagnosis (p<.05), treatment-associated reduction of depression symptoms (p<.001), improvement of delayed recall (p<.001), and baseline brain response to negative emotions (p<.05, whole brain corrected). Modeling the Age X Cortisol interaction suggested that for the participants below the median age of our sample, lower cortisol levels predicted a lower rate of MDD diagnosis, higher antidepressant effects and decreased brain reactivity in emotion regulation regions such as the anterior cingulate gyrus. On the contrary, in those above the median sample age, lower cortisol predicted a higher rate of MDD, less improvement in depression symptoms and memory performance, and more brain reactivity in the anterior cingulate.

**Conclusions:** Our results indicate that age moderates the relationship between peripheral cortisol levels and (1) MDD diagnosis, (2) brain reactivity to emotional stimuli, and (3) antidepressant-associated improvement in depression and memory symptoms. These results indicate that previous disparities in the literature linking peripheral cortisol levels with depression characteristics and treatment response may critically relate, at least in part, to the age of the patients studied.

## 1. INTRODUCTION

Because of the immense and growing cost of Major Depressive Disorder (MDD) to individuals (Ferrari et al., 2013) and society (Greenberg et al., 2015), uncovering physiological biomarkers associated with depression that may predict treatment response is of high importance. Data accumulated over four decades of research indicate that the hypothalamic-pituitary-adrenal (HPA) axis plays a key role in the pathophysiology of MDD (Stetler and Miller, 2011; Wolkowitz et al., 2009) and might have utility for prediction of depression symptom outcome to psychological treatments (Fischer et al., 2017). However, inconsistent results in the relationship between HPA axis markers such as cortisol and MDD suggest the presence of moderators such as age (Stetler and Miller, 2011).

Although elevations in circulating cortisol (O’Brien et al., 2004; Stetler and Miller, 2011; Vreeburg et al., 2009) or nonsuppression of cortisol levels with dexamethasone (Raison and Miller, 2003) (possibly indicating cortisol insensitivity) have often been found in MDD, many studies of older adults find no difference in overall cortisol levels in individuals with MDD relative to healthy controls--e.g. (Conrad et al., 2008; Fabian et al., 2001; Ferrari et al., 2001). Hypocortisolism was even detected in a significant proportion of elderly depressed patients (Bremmer et al., 2007; Oldehinkel et al., 2001). Other conditions characterized by high degrees of negative affect in elderly populations, including bereavement (Ong et al., 2011) and post-traumatic stress disorder (Yehuda et al., 1995), have also been associated with lower cortisol levels. In regards to the worst of depression outcomes – suicide – systematic meta-analysis found that age moderated the effects of cortisol on predicting suicidal behavior, with younger individuals with high cortisol and older ones with low cortisol at greater risk (O’Connor et al., 2016).

Age may also moderate the effects of glucocorticoids on memory function (Heffelfinger and Newcomer, 2001; Lupien et al., 2009). For example, in elderly rats, basal corticosterone levels correlated negatively with spatial memory performance, but this was not true of younger rats (Yau et al., 1995). Amitriptyline improved spatial memory in young rats and increased corticosteroid hippocampal gene expression, but these effects were not seen in older rats, suggesting some antidepressant mechanisms related to glucocorticoids may be moderated by age. In contrast to these findings in rodents, two studies in humans (with MDD status unspecified) suggested that higher cortisol levels have detrimental effects on memory in older participants. A longitudinal study found that aging individuals who exhibited a rise in cortisol levels over the course of five years exhibited delayed memory impairments relative to those with decreasing and currently moderate cortisol levels (Lupien et al., 1998); another longitudinal study found similar evidence for a negative effect of increasing cortisol levels on memory in women but not in men (Seeman et al., 1997).

We first undertook a retrospective study to understand the relationship between cortisol and age in predicting depression, treatment outcome and improvement in memory with antidepressant treatments. Based upon the published literature, and motivated by research that links suicidal behavior to younger individuals with higher cortisol and older individuals with lower cortisol (O’Connor et al., 2016), we predicted that age would moderate the effects of cortisol on depression treatment outcome, in that younger individuals with higher cortisol and older individuals with lower cortisol would improve less symptomatically and in regards to delayed memory function after a course of antidepressants. In a separate small sample of MDD patients, we prospectively sought to understand whether a central neurophysiological relationship might underlie these seemingly disparate peripheral cortisol phenotypes by conducting a pilot neuroimaging study of the relationship between cortisol and age in predicting response to negative emotional stimuli. We hypothesized that higher activity in the pregenual anterior cingulate gyrus, which previously has been correlated with antidepressant treatment resistance (Mayberg, 2009), is observed in younger individuals with higher cortisol and older individuals with lower cortisol.

## 2. Methods

### 2.1. Study participants

66 unmedicated MDD patients and 75 matched healthy controls (HC) were included in the retrospective analysis, and eight patients with MDD were enrolled prospectively in the neuroimaging study (Table 1). Participants shared the following recruitment and inclusion characteristics: They were recruited by flyers, bulletin board notices, newspaper ads, Craigslist postings, and clinical referrals. All procedures were approved by the Committee on Human Research of the University of California, San Francisco (UCSF) and the open label treatment study was registered on clinicaltrials.gov (NCT00285935). Since MRI studies were performed at the VA Medical Center in San Francisco, the corresponding procedures were also approved by the VA Medical Center in accordance with the Declaration of Helsinki. Study participants gave written informed consent to participate in this study and were compensated for participating. Those receiving antidepressant treatment were not charged.

**Table 1.** Demographic and Clinical Factors

|  | MDD base (N = 66) | HC (N = 75) | MDD treatment (N = 34) |
| --- | --- | --- | --- |
| Sex (% female) | 58 | 61 | 66 |
| Age (years) [range] | 38.8 ± 13.8 [19-68] | 37.3 ± 13.3 [20-69] | 37.4 ± 12.1 [19-65] |
| Race (%) |  |  |  |
| Caucasian | 77 | 84 | 77 |
| African American / Black | 12 | 10 | 11 |
| Asian | 3 | 3 | 3 |
| Other | 8 | 3 | 9 |
| Ethnicity |  |  |  |
| %Hispanic | 17 | 12 | 20 |
| Education (years) | 16.1 ± 2.2 | 16.7 ± 1.8 | 16.4 ± 2.2 |
| Age of MDD onset | 16.4 ± 8.7 | NA | 15.6 ± 9.2 |
| QIDS-SR baseline | 13.1 ± 3.8 | 1.9 ± 1.3 | 13.3 ± 3.3 |
| HAMD-17 baseline | 20.1 ± 3.3 | NA | 19.6 ± 3.0 |
| AM serum cortisol baseline (µg/dL) | 9.6 ± 4.6 | 9.7 ± 6.0 | 9.9 ± 4.7 |

HAMD-17=Hamilton Depression Rating Scale 17 item; HC=healthy control; MDD=Major Depressive Disorder; NA=not available; QIDS-SR=Quick Inventory of Depression Symptoms – Self Report; µg/dL=micrograms per deciliter.

Depressed participants were diagnosed with a current major depressive episode, without psychotic features, with the Structured Clinical Interview for DSM IV-TR Axis I Disorders (SCID) (First et al., 2015) and this diagnosis was confirmed by clinical interview with a board-certified psychiatrist. They scored ≥17 on the 17-item Hamilton Depression Rating Scale (HAMD) (Hamilton, 1960). Exclusion criteria for MDD patients were psychotic symptoms during their current major depressive episode, history of psychosis outside of a mood disorder episode, bipolar disorder, post-traumatic stress disorder, any eating disorder or severe trauma within one month of entering the study, and substance abuse or dependence (including alcohol) within six months of entering the study. Co-morbid anxiety disorders, with the exception of PTSD, were not exclusionary if MDD was considered the primary diagnosis. None of the study participants had acute illnesses or infections, chronic inflammatory disorders, neurological disorders, or any other major medical conditions considered to be potentially confounding (e.g., cancer, HIV, diabetes, history of cardiovascular disease or stroke, etc.), as assessed by history, physical examinations and routine blood screening. All study participants were required to be free of any psychotropic medications, hormone supplements, ACE-inhibitors, steroid-containing birth control or other potentially interfering medications, and had not had any vaccinations, for at least six weeks prior to enrollment in the study. None were taking vitamin supplements above the U.S. recommended daily allowances (e.g., 90 mg/day for Vitamin C). Short-acting sedative-hypnotics were allowed as needed up to a maximum of three times per week, but none within one week prior to participation. Prior to each study visit, all participants tested negative for drugs of abuse with a urine toxicology screen (marijuana, cocaine, amphetamines, phencyclidine, opiates, methamphetamine, tricyclic antidepressants, and barbiturates), and women of child-bearing potential tested negative for pregnancy with a urine screen. Individuals taking part in the imaging study were excluded for any MRI contraindications.

### 2.2. Blood draw preparations

Participants were admitted as outpatients to the UCSF Clinical and Translational Science Institute between 8 a.m. and 11 a.m., having fasted (except water) since 10 p.m. the night before. Participants were instructed to sit quietly and relax for 25 - 45 minutes before blood samples were obtained. Blood was collected into serum separator tubes, and serum was saved and frozen at −80 degrees C until assay. Cortisol assays for retrospective analysis were conducted in batches based on year of recruitment (2006–2008 and 2011– 2014) as previously described (Hough et al., 2017). For further details of cortisol assays, please refer to supplementary information.

### 2.3. Depression Symptom Measures

The severity of depressive symptoms was rated in depressed patients by self-report using the Quick Inventory of Depression Symptoms – Self Report (QIDS) (Rush et al., 2003), and by clinical observers using the HAMD, at baseline and (for treated patients) after 8 weeks of antidepressant medication. The QIDS was used as primary outcome because of its unidimensionality and high internal consistency (Reilly et al., 2015), and because it was normally distributed in our population whereas HAMD exhibited a leftward skew with most participants scoring at the cutoff point for study entry. Change in depression symptoms was calculated as post minus pre.

### 2.4. Cognitive Measures

The primary outcome studied was verbal and visual delayed recall due to previous work identifying a relationship between cortisol levels and delayed memory performance in aging (Lupien et al., 1998). The Hopkins Verbal Learning Task - Revised (HVLT) was used to assess verbal learning and memory (Benedict et al., 1998). Four different measures were calculated: the total immediate recall score, delayed recall after 20 minutes, recognition of cued words, and retention (proportion of words immediately recalled that were also remembered at the delay recall trial). The Brief Visuospatial Memory Test (BVMT) assessed visual learning and memory (Benedict et al., 1996). Three measures were calculated: the total immediate recall score, delayed recall after 20 minutes, and retention. To assess attention, WAIS-III forward and backward digit span tests were administered and the total score from both digit span trials was summed to provide an overall measure of performance (Wechsler, 1997). Raw scores for all measures were used. Not all participants completed cognitive measures: the HVLT was obtained on 27 MDD participants and the BVMT on 18 participants at follow up. All BVMT participants were drawn from the 2011-2014 pool comprising batch 2.

### 2.5. Antidepressant treatment

Based on prior medication history and side effect profile, participants were assigned open-label to receive an antidepressant medication from the class of selective serotonin reuptake inhibitors (SSRI). Participants were begun at a low dose which was titrated to therapeutic range according to standard clinical guidelines (American Psychiatric Association, 2010).

### 2.6. fMRI task

While undergoing MR imaging, MDD participants (median age 41, range 29-50) were shown images projected on a mirror of emotional faces (angry, fearful or neutral) or shapes (horizontal or vertical ovals, or circles) in a block design, as previously described (Hariri et al., 2002). 7 blocks were completed with a shape block followed by emotion block and ending with a shape block for a total of 4 blocks of shapes and 3 of faces. For full fMRI task details, please refer to supplementary information.

### 2.7. fMRI acquisition

MR Images were acquired on a 3 Tesla Siemens Skyra (Erlangen, Germany). For MRI acquisition parameters, please refer to supplementary information.

### 2.8. Statistical analysis

Statistical analysis was conducted with R version 3.2.3 (R Core Team, 2016). For predictive analyses of diagnosis and outcome, cortisol levels were log transformed and then converted to z scores by batch to facilitate comparison. Pearson correlations assessed the relationships among baseline variables with alpha=.05. Logistic regression was used to model the relationship between cortisol, age and MDD status. Linear models were used to predict treatment outcome. The data were examined for outliers and one participant was identified who scored more than 3 standard deviations below the mean on multiple cognitive tests at baseline (HVLT, BVMT, Digit Span), and this participant was removed from analysis of cognitive results. All model terms were presented as Beta±SE.

To model how the relationship between cortisol and clinical outcome was moderated by age, the results were plotted using a sliding window. Participants were first ordered from 1 to 34 on the basis of age. Sliding windows of 9 consecutive participants (e.g., participants 1 through 9, 2 through 10, 3 through 11, etc.) were obtained to cluster participants of similar age together. For each window of similarly aged participants, a linear regression model was performed using the relevant outcome as the dependent variable and cortisol as the independent variable. To facilitate visualization of the results, the beta values (for the slope of cortisol predicting the outcome variable in each sliding window) were plotted on the y axis versus the median of age of each sliding window on the x axis.

For the acquired fMRI data, analyses were conducted using AFNI (Cox, 1996). For full fMRI preprocessing details, please refer to supplementary information. Two general linear tests (GLT) were computed within participant: (angry + fearful) vs neutral, and faces vs shapes. All time points not accounted for by regressors constituted the baseline. A 3^rd^ order Legendre polynomial was included in the baseline to model slow signal drift.

To relate differences in processing stimuli type to cortisol and age, voxel-wise robust linear regression was performed in R. Robust regression is inherently more conservative and resistant to the presence of outliers in data (Huber, 1964) and as such has been recommended for use in neuroimaging data (Wager et al., 2005). Robust regression to predict brain activation (BA) was performed by calculating the difference in the contrast of threatening (angry + fearful) vs neutral faces and using this as the dependent variable, with the independent variable set as log cortisol, age and their interaction (cortisol scores did not need to be converted to z scores as in the clinical analysis because all samples were run in the same batch).

### 2.9. Correction for multiple comparisons

Task-based analysis and robust regressions were each required to pass voxel-wise two-tailed statistical thresholds (voxel-wise *p*<0.05; task analysis t(7)=2.36, regression t(4)=2.78). Voxels were further required to be part of a cluster of minimum volume, determined by a Monte-Carlo method, to control for multiple comparisons. The smoothing of the data was accounted for using the average of the spatial auto-correlation function (ACF) parameters from the individual participant’s data (Cox et al., 2017). Alpha was set at *p*<0.05. For the task-analysis, the minimum cluster sizes were 3,547μL, *p<*0.05. For the regression analyses, clusters were required to meet a minimum cluster size of 3,547μL, p<0.05.

## 3. RESULTS

### 3.1. Clinical sample

The clinical samples and matched HC were comparable on baseline demographic measures (Table 1). On average, they were mostly female, college educated, and Caucasian. Depression severity of MDD participants was on average moderate as assessed by both the HAMD and the QIDS, whereas the low score of HC on the QIDS was consistent with the absence of depression (these participants were not scored with the HAM-D17).

### 3.2. Prediction of MDD diagnosis

To predict MDD status, we used the overall MDD participant pool at baseline (termed “MDD base”) and HC samples (Table 1) and coded presence of MDD in a binary fashion. With logistic regression, neither Cortisol nor Age predicted MDD status (both p >.4). However, adding a Cortisol X Age interaction term into the model resulted in significance of the Cortisol term (B=1.37±0.64, p<.05) as well as the Cortisol X Age interaction (B=-0.04±0.02, p<.05), whereas Age was not a predictor (B=0.0±0.0, p=.4). To characterize what the Cortisol X Age term significance meant, we performed a median split on cortisol (z score = 0) and age (36 years). This yielded the following percentages of participants with MDD: low age and low cortisol 40%, low age and high cortisol 48%, high Age and low cortisol 56%, high Age and high Cortisol 42%. This suggested that the significance of the Cortisol X Age interaction was due to a modestly increased risk of MDD in those younger individuals with higher cortisol levels and older participants with lower cortisol levels.

### 3.3. Correlation among baseline factors, depressive symptoms and outcomes in MDD participants

In the sample of MDD participants treated with antidepressants (identified in Table 1 as “MDD treatment”, a subset of “MDD base”), we explored baseline correlations among variables and whether antidepressant-associated change in depression was associated with change in cognitive performance. QIDS and HAMD were significantly inter-correlated, (r=.54, p<.001). Age was significantly and negatively correlated with cortisol (r=-.35, p<.05) and baseline QIDS (r=-.51, p<.01) but not with HAMD (r=-.33, p=.06). Sex and BMI were not correlated with age or cortisol, nor with other baseline measures (p >.05 for all comparisons).

Improvement in self-rated (QIDS) and clinician-rated (HAMD) depression were significantly correlated (r=.58, p<.001). Reduction in QIDS over the 8-weeks of treatment was significantly associated with an improvement in HVLT delayed recall (r=-.41, p<.05) but reduction in HAMD was not (r=-.14, p >.4). Changes in depression scores were otherwise not associated with changes in HVLT or BVMT measures (p >.05 for all comparisons). Change in digit span was not associated with changes in depression or memory (p >.05 for all comparisons).

### 3.4. Effects of treatment

With treatment, self- and clinician-rated depression symptoms declined significantly as expected for a single group, open label treatment (Table 2). HVLT total learning score and delayed recall improved, but there were no changes in other verbal memory measures, visual learning or attention (Table 2).

**Table 2.** Depression and Cognition Changes After 8 Weeks Treatment

|  | pre | post | *t* (df) | p value |
| --- | --- | --- | --- | --- |
| QIDS-SR | 13.3 ± 3.3 | 7.21 ± 3.21 | 9.21 (33) | 10<sup>-9</sup> |
| HAM-D | 19.62 ± 3.04 | 9.71 ± 5.16 | 10.24 (33) | 10<sup>-10</sup> |
| HVLT Total | 27.42 ± 4.90 | 28.93 ± 3.71 | 2.44 (25) | 0.02 |
| HVLT Delayed Recall | 10.09 ± 2.05 | 10.81 ± 1.30 | 3.24 (25) | 0.003 |
| HVLT Recognition | 10.52 ± 3.33 | 11.19 ± 1.30 | 1.27 (24) | 0.22 |
| HVLT Retention | 96.94 ± 13.56 | 98.91 ± 10.37 | 1.21 (25) | 0.24 |
| BVMT Total | 26.79 ± 7.69 | 29.67 ± 3.45 | 1.27 (17) | 0.22 |
| BVMT Delayed Recall | 10.05 ± 2.82 | 10.89 ± .68 | .92 (17) | 0.37 |
| BVMT Retention | 82.34 ± 35.32 | 95.73 ± 5.25 | 1.23 (17) | 0.23 |
| Digit Span | 19.26 ± 4.72 | 19.67 ± 3.89 | .67 (32) | 0.51 |

BVMT=Binder Visual Memory Task; df=degrees of freedom; HAMD=Hamilton Depression Rating Scale 17 item; HC=healthy control; HVLT=Hopkins Verbal Learning Task; MDD=Major Depressive Disorder; *t*=paired t-test statistic; QIDS-SR=Quick Inventory of Depression Symptoms – Self Report

### 3.5. Age moderates effect of cortisol on predicting antidepressant treatment outcome (Fig 1 A, D)

**Figure 1.**
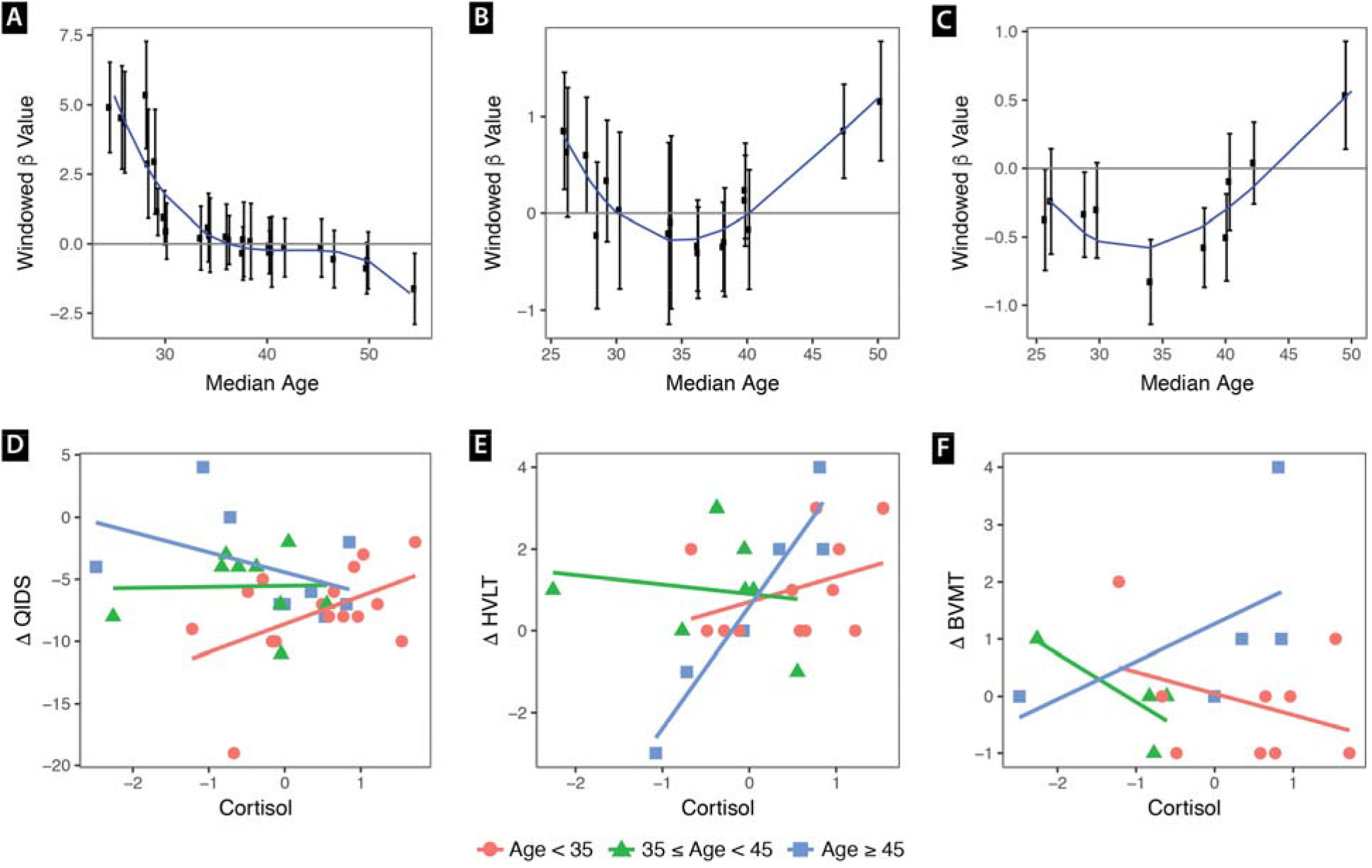
Legend. Panels A – C. Sliding window modeling of how cortisol predicted outcome in each age group. Participants were ordered on age. Successive windows iterated by one participant. Each window of 9 nine participants corresponds to one data point. Y axis shows Beta coefficient±one standard error of outcome change regressed on cortisol within each sliding window. X-axis indicates median age for the window of 9 participants in which the cortisol-outcome regression was performed. For panel A, negative Beta value indicates higher cortisol predicted improvement in depression symptoms. For panels B and C, positive Beta indicates that higher cortisol predicted improvement in verbal and visual delayed recall, respectively. A third order polynomial was used to model the curve. Panels D-F. Raw data points for cortisol (z scores) versus change in clinical outcome (pre – post). A negative delta for QIDS indicates clinical depression improvement, whereas a positive delta for HVLT and BVMT indicates memory improvement. Red circles show participants less than 35 years of age. Green triangles show participants between 35 and 44. Blue squares show participants 45 and older. The trend line was modeled within each age range. QIDS=Quick Inventory of Depression Symptoms – Self Report; BVMT=Binder Verbal Memory Test Delayed Recall; HVLT=Hopkins Verbal Learning Test Delayed Recall.

An interaction model predicted self-reported QIDS change from baseline to end of treatment with baseline Cortisol X Age (F(3,30)=7.85, adj R^2^=.38, p<.001). Within the model, all terms were significant predictors of QIDS change: Age (.19±.05, t=3.97, p<.001), Cortisol (7.44±.1.98, t=3.76, p<.001), and Cortisol X Age (−.18±.05, t=-3.72, p<.001). Inclusion of baseline QIDS or batch as a covariate did not substantially alter the results (Supplementary Materials). When tested individually for predicting change in QIDS, Age (.13±.05, F(1,32)=6.66, p=.015) and baseline QIDS (−.72±.16, F(1,32)=19.5, p<.001) were significant, but not Cortisol (−.24±.72, F(1,32)=.11, p=.74) or Batch (−1.46±1.34, F(1,32)=1.18, p=.29).

For clinically rated symptoms, an interaction model of Cortisol X Age predicted change in HAMD (F(4,29)=3.21, adj R^2^=.21, p<.05), with individual terms all reaching significance: Age (.20±.08, t=2.39, p=.02), Cortisol (8.36±3.37, t=2.49, p=.02), Cortisol X Age (−.17±.08, t=-2.11, p=.04). Adjusting for baseline HAMD did not reduce the overall significance of the model (F(4,29)=3.21, adj R^2^=.21, p<.05) but did lead to reduction of significance of the individual terms to trends: Age (6.69±3.38, t=1.98, p=.06), Cortisol (.14±.09, t=1.6, p=.12), baseline HAMD (−.57±.32, t=-1.80, p=.08), Cortisol X Age (−.13±.08, t=-1.60, p=.12). Adjusting for batch did not change the significance of individual terms but the addition of this term resulted in non-significance of the model (p=.09, Supplementary Info). When tested individually, neither Age (.11±.08, F(1,32)=1.76, p=.19), Cortisol (.90±1.03, F(1,32)=.77, p=.39), nor Batch (.75±1.97, F(1,32)=.15, p=.70), independently predicted change in HAMD, but baseline HAMD predicted improvement (−.78±.30, F(1,32)=6.96, p=.01).

### 3.6. Age moderates effect of cortisol on predicting improvement in delayed verbal recall (Fig 1 B, E)

Change in delayed verbal recall was predicted by an interaction model with Cortisol X Age (F(3,21)=4.01, adj R^2^=.27, p=.02), with individual terms of the model as follows: Age (−.02±.02, t=-.83, p=.41), Cortisol (−2.15±1.10, t=-1.95, p=.06), Cortisol X Age (.07±.03, t=2.70, p=.01). Additionally, adjusting for baseline delayed recall scores within the model led to a high prediction of variance (F(4,20)=18.61, adj R^2^=.75, p<10^−5^) and significance of all terms within the model: Age (−.04±.01, t=-2.95, p=.008), Cortisol (−1.45±.66, p=.04), baseline delayed recall (−.68±.11, t=-6.33, p<10^−5^), and Cortisol X Age (.04±.02, t=2.58, p=.02). Inclusion of sex, batch or change in digit span, QIDS or HAMD, in the model did not substantially reduce this significance (Supplementary Info).

The interaction of Cortisol X Age did not predict change in HVLT Total Recall, Recognition memory, or Retention (Supplementary Info). To determine whether lack of a Cortisol X Age effect was due to not accounting for initial performance, baseline scores of these variables were included in the model. This led to overall significance of the models, but the only terms significant were baseline scores (Supplementary Info), again reiterating lack of a Cortisol X Age effect on predicting these scores.

### 3.7. Effects of Cortisol X Age on change in delayed visual recall (Fig 1 C, F)

Because change in the delayed recall parameter for verbal memory (HVLT) was predicted by Cortisol X Age, we next tested whether Cortisol X Age predicted change in delayed visual recall using the BVMT. Although the interactive model including Age, Cortisol, and Cortisol X Age did not significantly predict delayed visual recall (p=.08, Supplementary Info), accounting for baseline performance led to significance of the model and the interactive effect of Cortisol X Age (F(4,13)=20.7, adj R^2^=.82, p<10^−4^). All terms in the model except age were significant: Age (−.02±.01, t=-2.56, p=.15), Cortisol (−1.16±.45, p=.02), baseline delayed recall (−.82±.12, t=-6.89, p<10^−4^), and Cortisol X Age (.03±.01, t=2.71, p=.02). Adjusting for sex or change in attention scores did not impact the significance of these findings, but adjusting for change in QIDS or HAMD resulted in reduction of significance of the Cortisol X Age interaction to trend level (p=.08, Supplementary Results).

### 3.8. Lack of Effect of Cortisol X Age on attention change

With respect to change in the digit span test, we found no predictive effect of age, cortisol, or their interaction on attention change (p > .5 for all independent variables). However, adding baseline digit span did result in significance of the model, with baseline digit span as the only significant predictor (Supplementary Results).

### 3.9. Effect of Cortisol X Age on brain response to negative emotional stimuli

In a prospective pilot study of eight participants with MDD, we studied brain responses to negatively valenced emotional faces. Comparing neural responses to faces and shapes revealed activation in two clusters: (1) fusiform face area (BA37), suggesting appropriately increased activation during facial processing, and (2) bilateral dorsolateral prefrontal cortex (suggesting increased cognitive control necessary for performing the matching while viewing faces, Figure 2A). To demonstrate an effect of negative emotions, we contrasted matching fearful and angry faces together versus neutral faces and performed a whole-brain analysis that demonstrated clusters of activation in ventromedial prefrontal cortex (vmPFC) and precuneus (Figure 2B).

**Figure 2.**
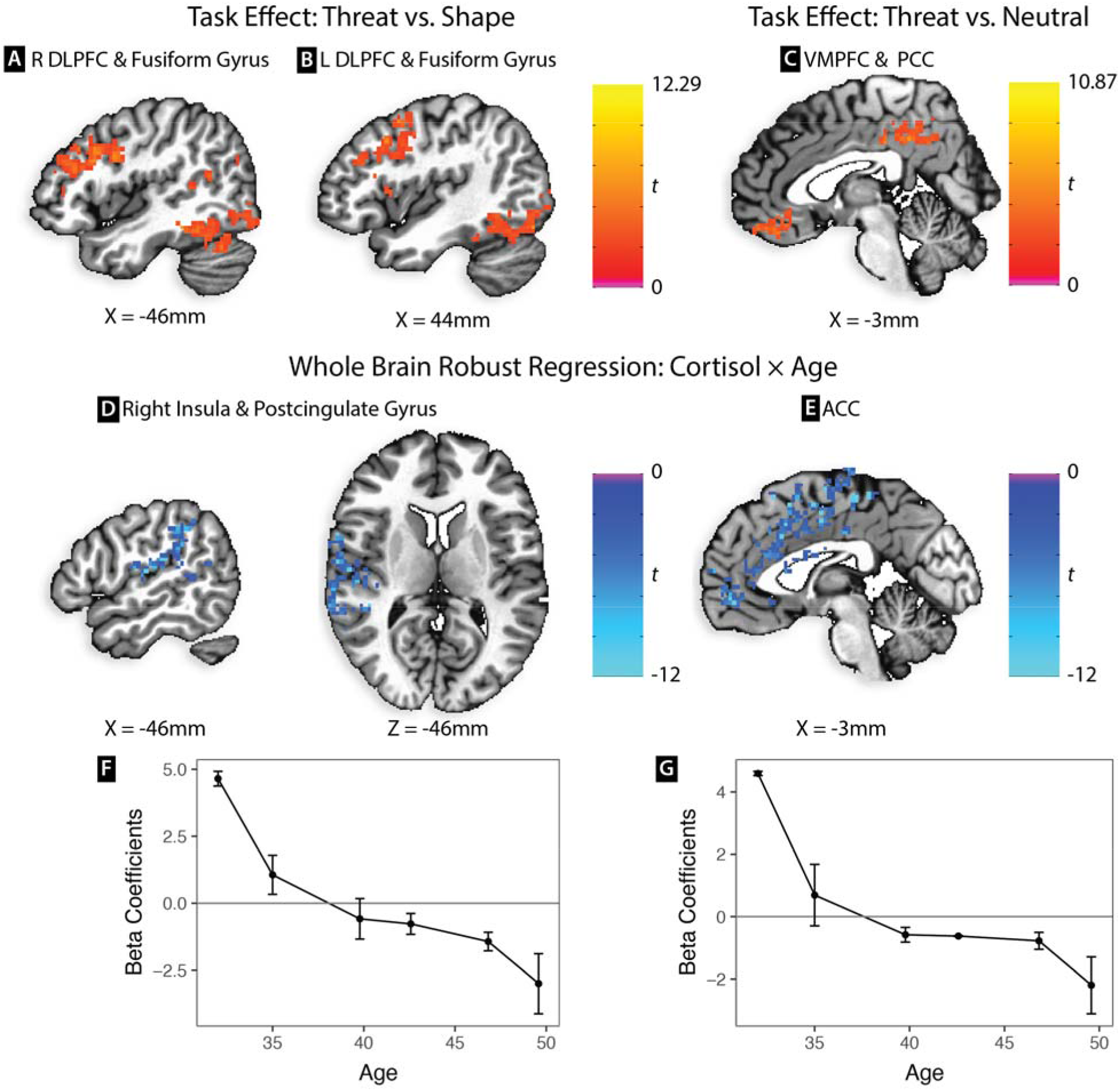
Legend. (A) and (B) Brain activation to negative emotion (angry and fearful) faces versus shapes. Labels indicate foci of activation clusters. (C) Task effect of brain activation to negative emotion faces compared to neutral faces. (D) sagittal and axial views and (E) sagittal view of clusters of activation predicted by Cortisol X Age using whole brain robust regression. (F) and (G) Sliding window modeling of how cortisol predicted brain activation in each age group, for clusters in (D) and (E), respectively. Y axis shows Beta coefficient±one standard error of brain activity regressed on cortisol within each sliding window of 3 participants. Successive windows iterated by one participant. X-axis indicates median age for the window of 3 participants in which the cortisol-outcome regression was performed. Positive Beta value indicates that higher cortisol predicted higher brain activation to negative emotional faces, and negative Beta value that higher cortisol predicted less brain activation. ACC=anterior cingulate cortex; DLPFC=dorsolateral prefrontal cortex; L=left; PCC=posterior cingulate cortex; R=right; X and Z coordinates shown in Right-Posterior-Inferior convention.

We then analyzed whether activation differences between emotional faces representing negative emotions (fearful and angry) vs neutral faces were predicted by Age, Cortisol, or Cortisol X Age, using robust regression with brain activation difference as the dependent variable. In this rather small sample, Age and Cortisol were not correlated (p >.6), and did not independently predict differential brain activation between threatening and neutral faces. However, large clusters of activation were predicted by the Cortisol X Age interaction, including right insula (BA41), bilateral supplementary motor area (BA6), and bilateral cingulate: subgenual (BA25), perigenual (BA24, BA23), dorsal anterior (BA24) and posterior (BA31, Figure 2C and 2D.) Modeling the results with a sliding window analysis (n=3 per window) suggested that younger individuals with higher cortisol and older individuals with lower cortisol had greater threat reactivity (Figure 2E), in a pattern reminiscent of self-reported QIDS change as a function of age (Figure 1A).

## 4. DISCUSSION

In a sample of MDD individuals who were physically healthy and unmedicated at study entry, age moderated the effects of cortisol on predicting several important outcomes: self- and clinician-rated symptomatic improvement with antidepressant treatment, and delayed recall with both verbal and visual stimuli. Cortisol, moderated by age, also weakly but significantly predicted MDD diagnostic status. For adults younger than the median age of our sample, modeling of the data indicated that higher levels of cortisol were associated with worse treatment response, while for participants older than the median, higher levels of cortisol were associated with better treatment response. For cognitive outcomes, there appeared to be little association with cortisol at lower ages but a strong association in the upper age range, most prominently in those over 45 years of age, in which relatively higher baseline levels of cortisol predicted improvement in both verbal and visual delayed memory. The improvements in memory and mood could neither be explained by covariates such as sex or baseline symptoms, nor by a commensurate change in attention. To our knowledge, this is the first study to find a consistent biomarker (cortisol level) effect predictive of MDD diagnostic status, memory improvement and reduction in depression symptoms, and to demonstrate its moderation by age.

Supporting these findings, in a prospective pilot neuroimaging study, cortisol demonstrated a different relationship to brain processing of negative emotions as a function of age. Higher levels of cortisol were associated with greater reactivity to negative emotional stimuli in the younger adult participants, but with lower reactivity to negative emotional stimuli in older adult participants. This was particularly evident in a large volume of anterior cingulate cortex, particularly the subgenual cingulate in which hyperactivity has also previously been associated with poor antidepressant treatment response (Dunlop et al., 2017; Mayberg, 2009), and the dorsal anterior cingulate which is involved in the experience of emotional pain (Eisenberger et al., 2003). Both brain regions serve as targets for neurosurgical procedures that reportedly ameliorate depression (Shields et al., 2008). The validity of our negative finding that basal cortisol alone was not associated with an increase in brain response to negative emotions is consistent with a report on 26 healthy participants that basal cortisol alone did not predict greater brain activation in response to negative emotional faces (Weldon et al., 2015). Taken together, our findings imply that simple linear models of circulating cortisol elevation across the entire age range may not be phenotypically valid in MDD.

Our non-imaging findings related to treatment outcome, though from a limited sample of 34 individuals, are consistent with two recent large meta-analyses that assessed the impact of age in moderating cortisol on prediction of outcome in varied clinical populations. First, O’Connor et al (2016) demonstrated a similar relationship between age and cortisol in predicting suicidal behavior. This meta-analysis analyzed 27 studies including over 2000 individuals and found an interactive effect between age and basal cortisol in predicting suicide attempt across patients with different diagnoses: those patients under 40 with higher cortisol and those over 40 with lower cortisol were at greatest risk. Interestingly, the inflection point for antidepressant treatment outcome in this study was also at about 40 years of age. The reason for this shift in midlife is unclear. Although the curve is roughly similar to that of fertility, inclusion of sex in our model did not reduce the significance of our findings or clearly suggest a sex-specific effect.

The finding that age moderated the effect of cortisol on predicting MDD diagnostic status is also broadly consistent with a meta-analysis of 26 studies containing over 5000 individuals that found a similar age X cortisol interaction to predict post-traumatic stress disorder (PTSD) risk (Morris et al., 2016). In this meta-analysis, younger individuals with higher cortisol and older individuals with lower cortisol in the period following a traumatic event were at greater risk for later development of PTSD. However, the inflection point in their study was approximately 30 years of age, perhaps suggesting that the physiological process underlying this finding might be accelerated in those with significant trauma reactivity, relative to those without PTSD (as comprised our sample).

The finding that higher cortisol levels in younger individuals was associated with higher MDD and worse treatment response is consistent with several prior reports. Stetler et al. (2011), for example, found that higher cortisol levels were evident with a moderate effect size in MDD using meta-analysis. Prior studies of cortisol as a predictor of MDD treatment with psychological (Fischer et al., 2017) and antidepressant treatment (Kin et al., 1997) have suggested that higher levels are a poor prognostic sign. To our knowledge, this is the first report to suggest that lower cortisol in older adults might be diagnostic and a poor prognostic indicator. Potential mechanisms of this effect are unclear but we suggest two possibilities related to changes in the HPA Axis with aging and stress: an overall reduction in HPA activity, or increased neural glucocorticoid receptor sensitivity which could result in lower peripheral cortisol levels via feedback inhibition. Both of these mechanisms have been postulated to underlie clinical symptoms often comorbid with depression, including somatic symptoms and pain syndromes, chronic fatigue, and stress related disorders, albeit irrespective of age (Fries et al., 2005; Heim et al., 2000; Raison and Miller, 2003).

The current study’s findings should be interpreted in light of its strengths and limitations. Strengths of our study included that our participants were medically healthy and unmedicated for a minimum of 6 weeks before baseline testing. Another strength is that our pilot neuroimaging results were obtained in an independent sample, but the results should be interpreted with caution given the low number of individuals imaged. Limitations included that we relied on a single basal morning cortisol level. Although variability of setting was mitigated by laboratory draw after a period of rest, the pulsatile nature of cortisol secretion could lead to variability of measurement across days. Second, our sample sizes for the retrospective clinical outcomes analysis and prospective neuroimaging were limited. In part to address this limitation, outcomes utilized continuous measures as opposed to splitting the sample into subgroups, and this also served to increase our power to find an effect. Participants were all younger than 70 years and our main findings may not apply to those over 70 years of age.

We conclude that the relationships of cortisol to MDD and antidepressant treatment outcome may be moderated by age, and this should be taken into account when studying their associations. Further research should sample cortisol at multiple times across several days in a home environment and in the lab, and it should involve larger samples across a broad range of ages. Using larger samples to study whether there may be a common neural signature underlying those with MDD who are younger with higher cortisol levels and older with lower cortisol levels may help to elucidate whether there is a common central neurophysiological process resulting in this biphasic peripheral relationship. Our findings suggest that combining peripheral and central biomarkers with age might help to link seemingly disparate hypercortisol vs hypocortisol findings reported in the literature on MDD. Further study of these mechanisms may help elucidate targets of antidepressant treatments.

## 5. CONTRIBUTORS

OMW, SHM, ESE, VIR, SM, DJM, TTY and FAJ were responsible for study concept and design. OMW, SHM, CMH, AM, VIR, DJM and FAJ contributed to the acquisition of data. FAJ and CGC performed initial data analysis and interpretation of findings. OMW, SHM, VIR, CMH, DL and SM assisted in interpretation of findings. FAJ and CGC drafted the manuscript. All authors critically reviewed content and approved the final manuscript.

## 6. ACKNOWLEDGEMENTS

Data acquisition was funded by grants from the National Institute of Mental Health (Grant Number R01 – MH083784). This project was also supported by National Institutes of Health/National Center for Research Resources (NIH/NCRR) and the National Center for Advancing Translational Sciences, National Institutes of Health, through UCSF-CTSI Grant Number UL1 RR024131. The University of Virginia Center for Research in Reproduction Ligand Assay and Analysis Core is supported by the Eunice Kennedy Shriver NICHD/NIH (NCTRI) Grant P50-HD28934. Author TTY was supported by the Brain and Behavior Research Foundation (formerly NARSAD) and grant number R01MH085734-05 from the National Institute of Mental Health. Author FJ was supported by grant #R21AG051970 from the National Institute on Aging. Additional financial support was obtained from the Tinberg Family and the O’Shaughnessy Foundation.

The authors gratefully acknowledge statistical consultation by Kevin Delucchi, PhD, the nursing and other staff of the UCSF CTSI Clinical Research Center, administrative assistance of Brenton Nier and Mina Cheema, and fMRI acquisition assistance from Donna Murray, PhD. None of the granting or funding agencies had a role in the design and conduct of the study; collection, management, analysis and interpretation of the data; and preparation, review, or approval of the manuscript. The Co-Principal Investigators, OMW, ESE, and SHM, had full access to all data in the study and take responsibility for the integrity of the data and the accuracy of the data analysis. The contents of this publication are solely the responsibility of the authors and do not necessarily represent the official views of the NIH.

